# Adaptive potential of a drug-targeted viral protein as a function of environmental stress

**DOI:** 10.1101/078428

**Authors:** Lei Dai, Yushen Du, Hangfei Qi, Christian D. Huber, Nicholas C. Wu, Ergang Wang, James O. Lloyd-Smith, Ren Sun

**Author notes:** These authors contributed equally to this work (Lei Dai, Yushen Du). Corresponding author: Lei Dai, Ph.D., Ren Sun, Ph.D.

## Abstract

RNA viruses are notorious for their ability to evolve rapidly under selection in novel environments. It is known that the high mutation rate of RNA viruses can generate huge genetic diversity to facilitate viral adaptation. However, less attention has been paid to the underlying fitness landscape that represents the selection forces on viral genomes. Here we systematically quantified the distribution of fitness effects (DFE) of about 1,600 single amino acid substitutions in the drug-targeted region of NS5A protein of Hepatitis C Virus (HCV). We found that the majority of non-synonymous substitutions incur large fitness costs, suggesting that NS5A protein is highly optimized in natural conditions. We characterized the adaptive potential of HCV by subjecting the mutant viruses to selection by the antiviral drug Daclatasvir. Both the selection coefficient and the number of beneficial mutations are found to increase with the level of environmental stress, which is modulated by the concentration of Daclatasvir. The changes in the spectrum of beneficial mutations in NS5A protein can be explained by a pharmacodynamics model describing viral fitness as a function of drug concentration. We test theoretical predictions regarding the distribution of beneficial fitness effects of mutations. We also interpret the data in the context of Fisher’s Geometric Model and find an increased distance to optimum as a function of environmental stress. Finally, we show that replication fitness of viruses is correlated with the pattern of sequence conservation in nature and viral evolution is constrained by the need to maintain protein stability.

## Introduction

In our evolutionary battles with microbial pathogens, RNA viruses are among the most formidable foes. HIV-1 and Hepatitis C Virus acquire drug resistance in patients under antiviral therapy. Influenza and Ebola virus cross the species barrier to infect human hosts. Understanding the evolution of RNA viruses is therefore of paramount importance for developing antivirals and vaccines and assessing the risk of future emergence events (Goldberg *et al.* 2012; Domingo *et al.* 2012; Metcalf *et al.* 2015). Comprehensive characterization of viral fitness landscapes, and the principles underpinning them, will provide us with a map of evolutionary pathways accessible to viruses and guide our design of effective strategies to limit antiviral resistance, immune escape and cross-species transmission (Turner and Elena 2000; Ke *et al.* 2015; Barton *et al.* 2016).

Although the concept of fitness landscapes has been around for a long time (Wright 1932), we still know little about their properties in real biological systems. Previous empirical studies of fitness landscapes have been constrained by very limited sampling of sequence space. In a typical study, mutants are generated by site-directed mutagenesis and assayed for growth rate individually. We and others have recently developed a high-throughput technique, often referred to as “deep mutational scanning” or “quantitative high-resolution genetics”, to profile the fitness effect of mutations by integrating deep sequencing with selection experiments in vitro or in vivo (Hietpas *et al.* 2011; Wu *et al.* 2013; Thyagarajan and Bloom 2014; Qi *et al.* 2014; Fowler and Fields 2014). This novel application of next generation sequencing has raised an exciting prospect of large-scale fitness measurements (Olson *et al.* 2014; Puchta *et al.* 2015; Li *et al.* 2016; Wu *et al.* 2016) and a revolution in our understanding of molecular evolution (He and Liu 2016).

The distribution of fitness effects (DFE) of mutations is a fundamental entity in genetics and reveals the local structure of a fitness landscape (Burch and Chao 2000; Eyre-Walker and Keightley 2007; Hietpas *et al.* 2011; Desai 2013; Jacquier *et al.* 2013; Bataillon and Bailey 2014; Chevereau *et al.* 2015; Bank *et al.* 2015). Deleterious mutations are usually abundant and impose severe constraints on the accessibility of fitness landscapes. In contrast, beneficial mutations are rare and provide the raw materials of adaptation. Quantifying the DFE of viruses is crucial for understanding how these pathogens evolve to acquire drug resistance and surmount other evolutionary challenges.

Most empirical studies of the DFE have been performed in a single, static environment (Eyre-Walker and Keightley 2007; Bataillon and Bailey 2014). A central challenge is to characterize the DFE, and its determinants, in fluctuating or heterogeneous environments where evolution typically occurs (e.g. fluctuating drug concentrations or a gradient across space). Previous studies on yeast have investigated the change in DFE across different levels of temperature and salinity (Hietpas *et al.* 2013; Bank *et al.* 2014). For bacteria, the fitness effects of mutations at different drug concentrations have been studied (Firnberg *et al.* 2014). One recent study has demonstrated that drug concentration modulates the shape of the DFE and determines the evolvability under new environments (Stiffler et al. 2015). In another study, the implications of differing drug concentrations on the adaptive landscape have been examined in the context of resistance evolution (Ogbunugafor *et al.* 2016). For viruses, the fitness effects of mutations have been measured across different hosts (Lalić *et al.* 2011; Vale *et al.* 2012). The shape of DFE of viruses has been inferred from experimentally passaged populations (Foll *et al.* 2014) and from patient data (Renzette *et al.* 2017).

In this study, we profile the DFE of ~1,600 single amino acid substitutions in a drug-targeted viral protein by combining selection experiment of a mutant library and deep sequencing. We examine the changes in DFE under varying levels of environmental stress by tuning the concentration of an antiviral drug. We test theoretical predictions regarding the distribution of beneficial fitness effects of mutations (Orr 2003). We also interpret the data in the context of Fisher’s Geometric Model (Martin and Lenormand 2006b) and find an increased distance to optimum as a function of environmental stress. Finally, we show that replication fitness of viruses is correlated with the pattern of sequence conservation in nature and viral evolution is constrained by the need to maintain protein stability.

## Results

### Profiling the fitness landscape of the drug-interacting domain of HCV NS5A protein

The system used in our study is Hepatitis C Virus (HCV), a positive sense single-stranded RNA virus with a genome of ~9.6 kb. HCV has been studied extensively in the past two decades in patients and in laboratory and provides an excellent model system to study viral evolution. We applied high-throughput assays to map the fitness effects of all single amino acid substitutions in domain IA (amino acid 18-103) of HCV NS5A protein (Methods). This domain is the target of several directly-acting antiviral drugs, including the potent HCV NS5A inhibitor Daclatasvir (DCV) (Gao *et al.* 2010).

To study the DFE of mutations of HCV NS5A protein, we conducted new selection experiments using a previously constructed saturation mutagenesis library of mutant viruses (Qi *et al.* 2014). Briefly, each codon in the mutated region was randomized to cover all possible single amino acid substitutions. We observed 2520 non-synonymous mutations in the plasmid library, as well as 105 synonymous mutations. After transfection to reconstitute mutant viruses, we performed selection in an HCV cell culture system (Lindenbach *et al.* 2005; Wakita *et al.* 2005). The relative fitness of a mutant virus to the wild-type virus was calculated based on the changes in frequency of the mutant virus and the wild-type virus after one round of selection in cell culture (Supplementary Figure 1). In our selection experiment, we grew 5 small sub-libraries (~500 mutants each) separately to reduce the noise in fitness measurements (Methods). The fitness data reported in this study is highly correlated to an independent experiment using the same plasmid library (Supplementary Figure 2) (Qi *et al.* 2014).

Our experiment provides a comprehensive profiling of the fitness effect of single amino acid substitutions (1565 out of 1634 possible substitutions, after filtering out low frequency mutants in the plasmid library). We grouped together non-synonymous mutations leading to the same amino acid substitution. As expected, the fitness effects of synonymous mutations were nearly neutral, while most non-synonymous mutations were deleterious (Figure 1). We found that the majority of single amino acid mutations had fitness costs and more than half of them were found to be significantly deleterious, or “lethal” (Methods). The fraction of lethal mutations (not shown explicitly in Figure 1) is 57.0% (932/1634) for single amino acid substitutions, 1.0% (1/105) for synonymous mutations and 90.6% (77/85) for nonsense mutations. The low tolerance of non-synonymous mutations in HCV NS5A, which is an essential protein for viral replication, is consistent with previous small-scale mutagenesis studies of RNA viruses (Sanjuan *et al.* 2004). Our data support the view that RNA viruses are very sensitive to the effect of deleterious mutations, possibly due to the compactness of their genomes (Elena *et al.* 2006; Rihn *et al.* 2013).

**Figure 1.**
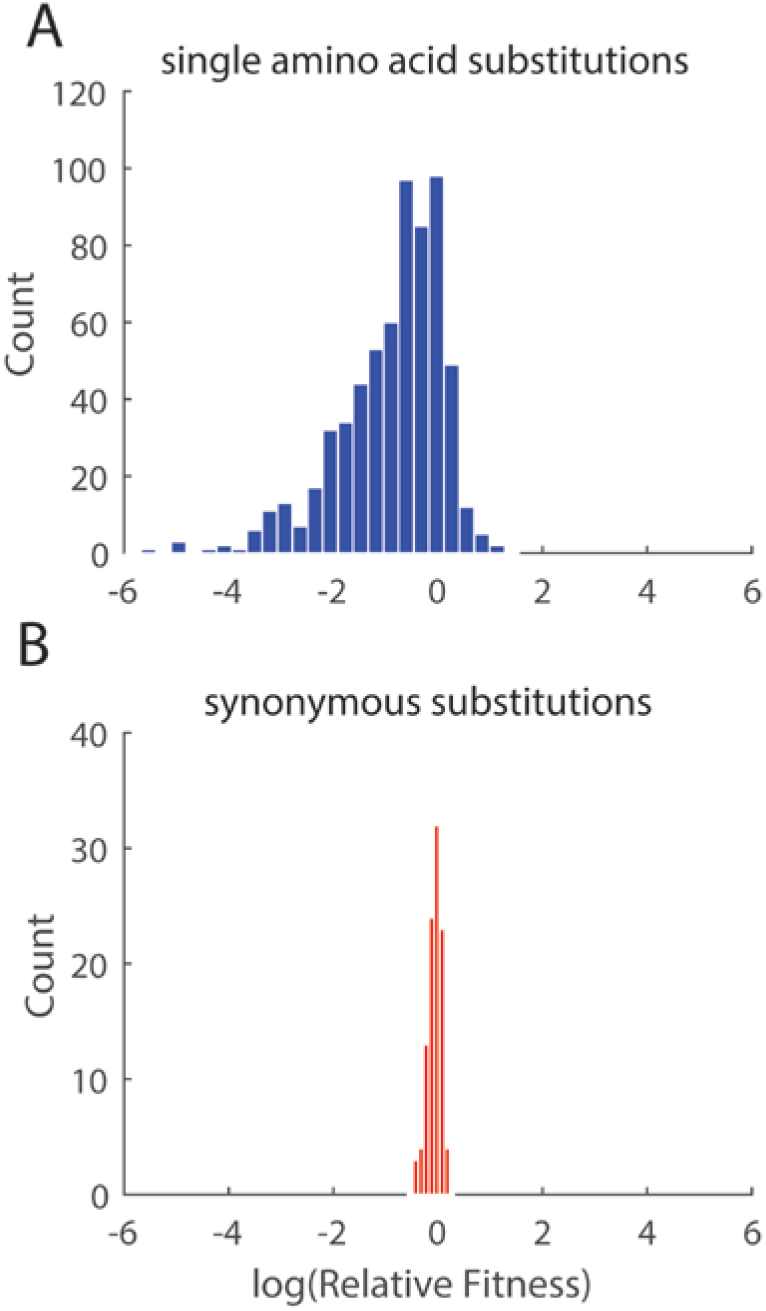
Distribution of fitness effects (DFE) of single amino acid substitutions in domain IA of HCV NS5A protein without drug selection. DFE of single amino acid substitutions (A) and synonymous substitutions (B). Lethal mutations are not shown in the histogram.

Using the distribution of fitness effects of synonymous mutations as a benchmark for neutrality, we identified that only 2.3% (37/1634) of single amino acid mutations are beneficial (Methods). The estimated fraction of beneficial mutations is consistent with previous small-scale mutagenesis studies in viruses including bacteriophages, vesicular stomatitis virus, etc. (Sanjuan *et al.* 2004; Burch *et al.* 2007; Silander *et al.* 2007; Eyre-Walker and Keightley 2007). Our results indicate that HCV NS5A protein is under strong purifying selection, suggesting that viral proteins are highly optimized in their natural conditions.

### Adaptive potential as a function of environmental stress

Beneficial mutations are the raw materials of protein adaptation (Eyre-Walker and Keightley 2007). In this study, we aimed to study the role of environmental stress in modulating the adaptive potential of drug-targeted viral proteins. In an independent study (Qi *et al.* 2014), the mutant library of HCV NS5A protein was selected under a single drug concentration ([DCV]=20 pM) to profile the effects of mutations on drug resistance. In this study, we selected the mutant library at 10, 40 and 100 pM of DCV. The drug concentrations were chosen based on in vitro IC_50_ of wild type HCV virus (~20 pM) to represent different levels of environmental stress (mild, intermediate and strong).

By tuning the concentration of DCV, we observed a change in the DFE (Supplementary Table 1&2), particularly of beneficial mutations (Figure 2A). At higher drug concentrations, we observed an increase in the median selection coefficient (Figure 2B) as well as the total number of beneficial mutations (Figure 2C, Supplementary Table 3). We further tested whether the shape of this distribution changed under drug selection. Previous empirical studies supported the hypothesis that the DFE of beneficial mutations is exponential (Orr 1998, 2003, 2006; Imhof and Schlötterer 2001; Sanjuan *et al.* 2004; Rokyta *et al.* 2005; Cowperthwaite *et al.* 2005; Kassen and Bataillon 2006; Burch *et al.* 2007; Carrasco *et al.* 2007; MacLean and Buckling 2009; Peris *et al.* 2010; Bataillon *et al.* 2011). Following a maximum likelihood approach, we fit the DFE of beneficial mutations to the Generalized Pareto Distribution (Supplementary Figure 3, Methods). The fitted distribution is described by two parameters: a scale parameter (**τ**), and a shape parameter (**κ**) that determines the behavior of the distribution’s tail. Using a likelihood-ratio test (Beisel *et al.* 2007), we found that our data are consistent with the null hypothesis that the DFE of beneficial mutations is exponential(κ = 0) (Supplementary Table 4).

**Figure 2.**
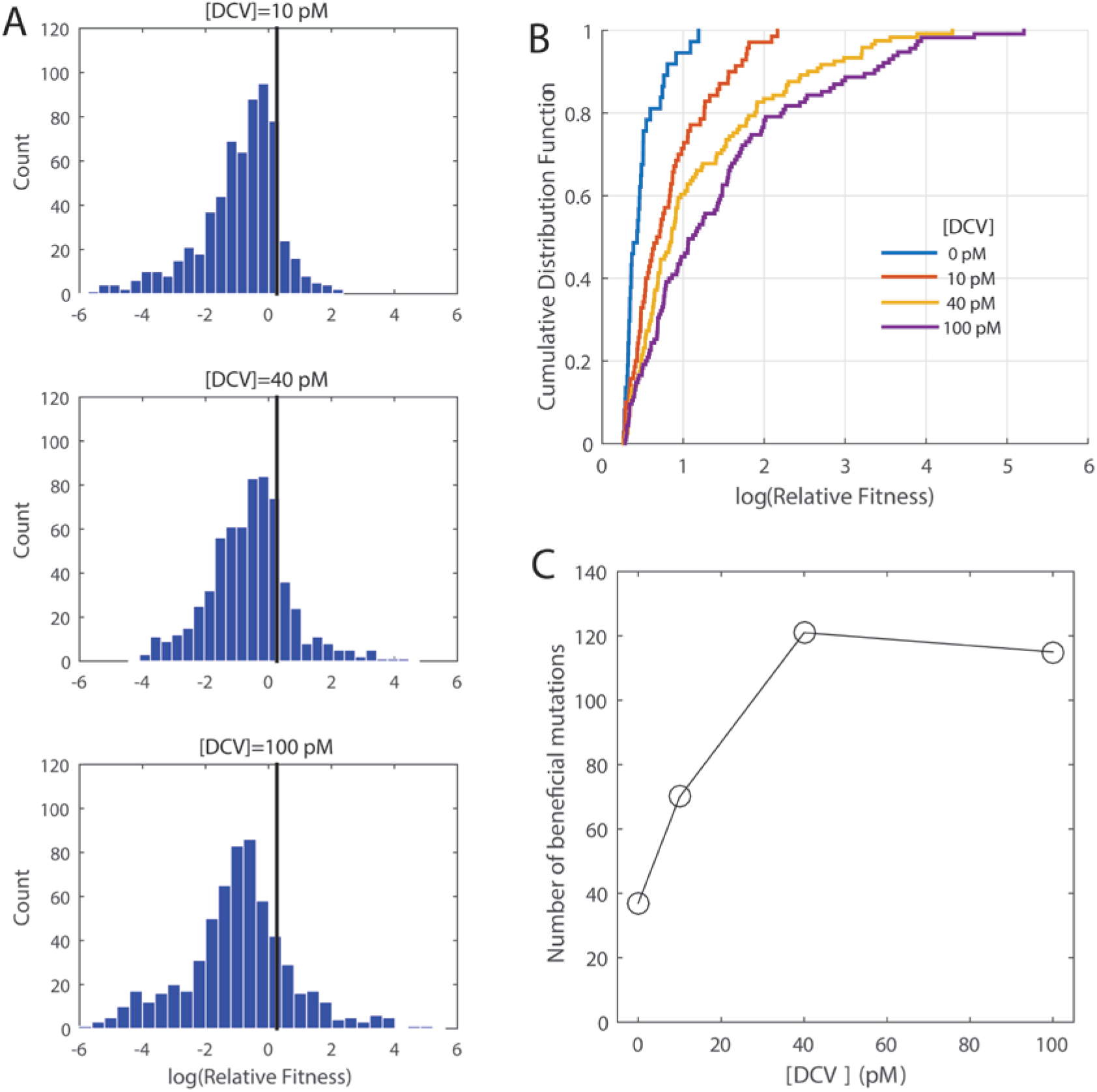
The spectrum of beneficial mutations changes under increasing environmental stress imposed by the antiviral drug Daclatasvir. (A) DFE of single amino acid substitutions in domain IA of HCV NS5A protein under increasing environmental stress by Daclatasvir. The black line indicates the threshold used for classifying beneficial mutations (Methods). (B) The cumulative distribution function of the fitness effect of beneficial mutations. (C) The number of beneficial mutations as a function of environmental stress imposed by Daclatasvir.

Furthermore, we used a maximum-likelihood approach to fit a displaced-gamma distribution to the DFE to estimate the distance to the phenotypic optimum in Fisher’s Geometric Model (FGM) (Martin and Lenormand 2006b; Bank *et al.* 2014) (Supplementary Figure 4). The displaced-gamma distribution has the shape of a negative gamma distribution, shifted by a parameter *s*_0_ that indicates the distance of the initial genotype (i.e. wild-type) to the optimum (Methods). Estimated distances to the optimum under different conditions are summarized in Supplementary Table 5. In accordance with theoretical expectations, we found that the distance to the optimum increased as the level of environmental stress increased (i.e. increasing drug concentration).

### The effects of mutations on drug resistance and replication fitness

Our results show that the adaptive potential of proteins is modulated by the strength of environmental stress. The changing spectra of beneficial mutations upon drug treatment can be explained by a pharmacodynamics model describing viral fitness as a function of drug concentration (i.e. phenotype-fitness mapping) (Figure 3A).

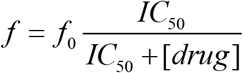

where *f*_0_ is the fitness without drug selection and *IC*_50_ is the half inhibitory concentration. The absolute fitness *f* decreases with drug concentration[*drug*]. In this paper, we define a drug-resistant mutant as any viral variant that is less inhibited than the wild type for some drug concentration, i.e. higher *IC*_50_ than wild-type (Rosenbloom *et al.* 2012).

**Figure 3.**
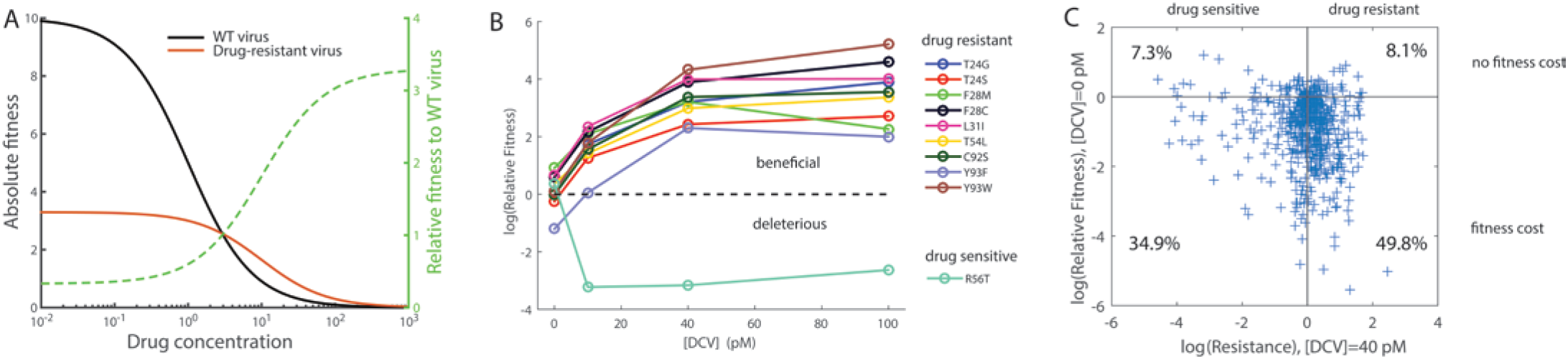
The adaptive potential under drug selection is determined by the effects of mutations on replication fitness and drug resistance. (A) Hypothetical dose response curves of the wild-type virus and a drug-resistant mutant virus. The absolute fitness *f* decreases with drug concentration [*drug*] 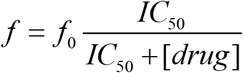, where *f*_0_ is the fitness without drug selection and *IC*_50_ is the half inhibitory concentration. Compared to the wild-type virus, the hypothetical drug-resistant mutant carries a fitness cost (smaller *f*_0_) but is less sensitive to drug inhibition (larger *IC*_50_). Relative fitness of the drug-resistant mutant is expected to increase with drug concentration. (B) Relative fitness of validated drug-resistant and drug-sensitive mutants (Supplementary Figure 5) as a function of [DCV]. (C) The effects of mutations on replication fitness (i.e. fitness without drug) and drug resistance score W at [DCV]=40 pM (Methods).

Mutations that reduce a protein’s binding affinity to drug molecules (i.e. less inhibited by the drug) may come with a fitness cost (i.e. smaller *f*_0_ than wild-type). Thus, a drug-resistant mutant that is deleterious in the absence of drug may become beneficial under drug selection, leading to an increase in the number of beneficial mutations. Moreover, the relative fitness of the drug-resistant mutant is expected to increase with stronger selection pressure (Figure 3A, dashed line). The dose response curves were previously measured for a set of mutants constructed by site-directed mutagenesis (Supplementary Figure 5) (Qi *et al.* 2014). Indeed, we found that the relative fitness of validated drug-resistant mutants increased at higher drug concentration (Figure 3B); in contrast, drug-sensitive mutants became less fit under drug selection.

Furthermore, we showed that the effects of mutations on drug resistance can be estimated from the fitness data and the results were generally consistent with estimates based on the dose response curves (Supplementary Figure 6, Methods). Among all the non-lethal single amino acid substitutions profiled in our HCV NS5A protein library, we found that roughly half of the mutations increased resistance to DCV (i.e. improved new function) at the expense of replication fitness without drug (Figure 3C, Spearman’s ρ= -0.13, p=8.3×10^-4^). This group of resistance mutations (lower right section in Figure 3C) can become beneficial when the positive selection imposed by the antiviral drug is strong, leading to an increase in the supply of beneficial mutations at higher drug concentrations. We found no association between drug resistance and fitness cost (Fisher’s exact test, p=0.26), suggesting that there is no or very weak tradeoff in adaptation of NS5A protein under the two different environments (i.e. with and without DCV selection).

### Deleterious mutations as evolutionary constraints

While beneficial mutations open up adaptive pathways to genotypes with higher fitness, mutations that severely reduce replication fitness impose constraints on the evolution of viruses and are less likely to contribute to adaptation through gain of function. We analyzed sequence diversity of HCV sequences identified in patients from the HCV sequence database of Los Alamos National Lab (Methods). As expected, we found that amino acid sites with high fitness costs are often highly conserved (Figure 4A). The sequence diversity at each site was highly correlated to the replication fitness (the median fitness of observed mutants at each site) measured in our study (Spearman’s ρ=0.82, p=1.8×10^-21^).

**Figure 4.**
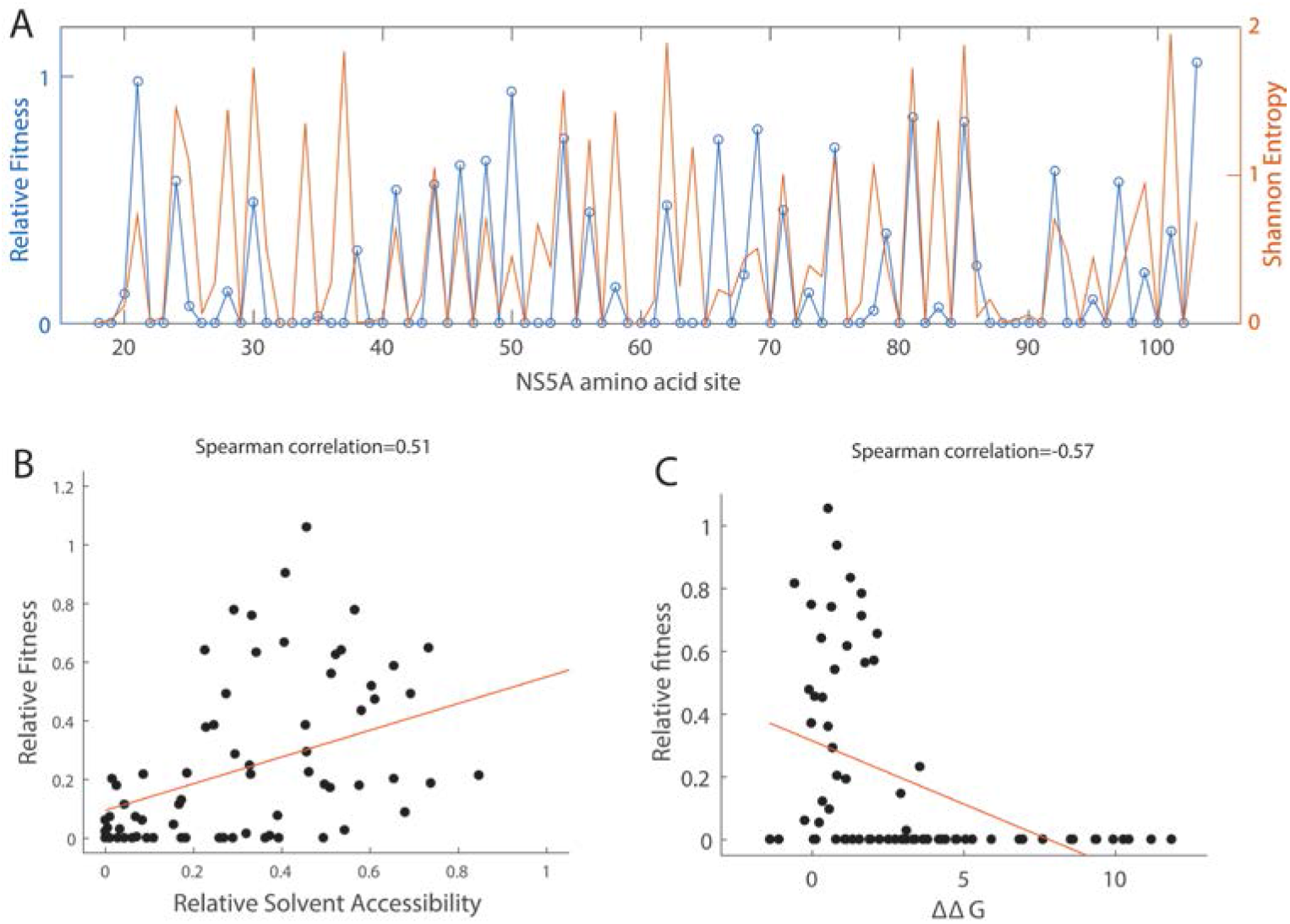
Mutations with deleterious fitness effects reveal constraints of protein evolution. (A) The pattern of sequence conservation observed in patient sequences is highly correlated to the replication fitness measured in cell culture. (B) Mutations at amino acid sites with lower solvent accessibility tend to incur larger fitness costs. (C) Mutations at amino acid sites with larger effects on destabilizing protein stability (ΔΔG>0) tend to reduce the viral replication fitness. Changes in folding free energy ΔΔG (Rosetta Energy Unit) of NS5A monomer were predicted by PyRosetta. The median ΔΔG at each amino acid site is shown. In (A-C), the median fitness of observed mutants at each amino acid site is shown. In (B) and (C), red lines represent the fits by linear regression and are only used to guide the eye.

To understand the biophysical basis of mutational effects (Liberles *et al.* 2012), we took advantage of the available structural information (Supplementary Figure 7A). The crystal structure of NS5A domain I is available excluding the amphipathic helix at N-terminus (Tellinghuisen *et al.* 2005; Love *et al.* 2009). We found that the fitness effects of deleterious mutations at buried sites (i.e. with lower solvent accessibility) were more pronounced than those at surface exposed sites (Figure 4B, Spearman’s ρ=0.51, p=5.1×10^-6^; Supplementary Figure 8A) (Ramsey *et al.* 2011). Moreover, we performed simulations of protein stability for individual mutants using PyRosetta (Methods) (Das and Baker 2008; Chaudhury *et al.* 2010). A mutation with ΔΔG>0, i.e. shifting the free energy difference to favor the unfolded state, is expected to destabilize the protein. We found that mutations that decreased protein stability led to reduced viral fitness (Figure 4C, Spearman’s ρ= -0.57, p=1.5×10^-7^). For example, mutations at a stretch of highly conserved residues (F88-N91) that run through the core of NS5A protein tended to destabilize the protein and significantly reduced the viral fitness. Mutations that increase ΔΔG beyond a threshold (~5 Rosetta Energy Unit) were mostly lethal (Supplementary Figure 8B). This is consistent with the threshold robustness model, which predicts that proteins become unfolded after using up the stability margin (Bloom *et al.* 2005; Wylie and Shakhnovich 2011; Olson *et al.* 2014). Also, we note that mutations can be deleterious because they impair protein function rather than destabilize the protein, so the correlation between protein stability and fitness is not expected to be perfect. The level of correlation between ΔΔG and fitness that we observed is similar to previous studies in other proteins (Firnberg *et al.* 2014; Wu *et al.* 2015).

## Discussion

Site-directed mutagenesis and experimental evolution are traditional approaches to examine the DFE (Domingo-Calap *et al.* 2009; Sanjuán 2010; Levy *et al.* 2015; Visher *et al.* 2016). Both methods provide pivotal insights into the shape of the DFE, yet with limitations. The site-directed mutagenesis approach requires fitness assays for each individual mutant and can only provide a sparse sampling of mutations. In experimental evolution, the sampling of sequence space via de novo mutations is biased towards large-effect beneficial mutations, as they are more likely to fix in the population. In contrast, the deep mutational scanning approach (Fowler and Fields 2014), which utilizes high-throughput sequencing to simultaneously assay the fitness or phenotype of a library of mutants, allows for unbiased and large-scale sampling of fitness landscapes and thus is ideal for studying the characteristics of empirical DFE. The downside of this high-throughput approach is that the fitness measurements can be noisy, especially for large mutant libraries (Matuszewski *et al.* 2016). In our experiment, we divided the mutant library into smaller sub-libraries (~500 mutants) in selection experiments. We compared the data to an independent experiment and found that the fitness estimates were largely reproducible (Supplementary Figure 2). We also showed that the observed change in the DFE under different conditions was consistent with validation experiments (Figure 3). Since this study is focused on the properties of the entire distribution of mutations rather than the effects of specific mutations, our findings on the general patterns of DFE are robust to the errors in fitness estimates. Our study quantified the fitness effects of single amino acid substitutions in the drug-targeted region of an essential viral protein. In general, the empirical DFE of HCV NS5A was consistent with previous findings that viral proteins were highly optimized in the natural condition and very sensitive to the effects of deleterious mutations.

One crucial but often overlooked point is that DFE will vary as a function of the environment (Martin and Lenormand 2006a; Lalić *et al.* 2011; Stiffler *et al.* 2015). In the study by Stiffler *et al.* 2015, the level of environmental stress is controlled by ampicillin concentration. Because TEM-1’s function is to degrade ampicillin, deleterious mutations that impair the enzyme function (“loss-of-function”) would become more deleterious at higher dose of ampicillin. In our system, we do not expect the dose of Daclatasvir to alter the strength of purifying selection on maintaining HCV NS5A protein’s function in viral replication. Indeed, we do not find much difference on the deleterious side of DFE across different environments. Instead, we have observed significant changes on the beneficial side of DFE as a function of the drug dose. Because HCV NS5A protein is not well adapted in the novel environment of Daclatasvir selection, the effect of drug resistance mutations (“gain-of-function”) becomes more beneficial at higher drug dose. Moreover, due to the pleiotropic effect of mutations on drug resistance and replication fitness (Figure 3), there is an increasing supply of beneficial mutations at higher drug dose.

Although different systems have distinct protein-drug interactions that lead to different resistance profiles (Robinson *et al.* 2011), the results in our study provide a general framework to study DFE of drug-targeted proteins. Future studies along this line will further our understanding of how proteins evolve new functions under the constraint of maintaining their original function (Soskine and Tawfik 2010), as exemplified in the evolution of resistance to directly-acting antiviral drugs (Rosenbloom *et al.* 2012). Quantifying the characteristics of DFE of drug-targeted proteins under different environments (e.g. varying levels of environmental stress, or conflicting selection pressures), would allow us to assess repeatability in the outcomes of viral evolution (de Visser and Krug 2014) and guide the design of therapies to minimize drug resistance (Ogbunugafor *et al.* 2016).

## Conclusions

Many viruses adapt rapidly to novel selection pressures, such as antiviral drugs. Understanding how pathogens evolve under drug selection is critical for the success of antiviral therapy against human pathogens. By combining deep sequencing with selection experiments in cell culture, we have quantified the distribution of fitness effects of mutations in the drug-targeted domain of Hepatitis C Virus NS5A protein. Our results indicate that the majority of single amino acid substitutions in NS5A protein incur large fitness costs. By subjecting the mutant viruses to selection under an antiviral drug, we find that the adaptive potential of viral proteins in a novel environment is modulated by the level of environmental stress. We test theoretical predictions regarding the distribution of fitness effects of mutations. Finally, we show that viral evolution is constrained by the need to maintain protein stability.

## Materials and Methods

### Mutagenesis

The mutant library of HCV NS5A protein domain IA (86 amino acids) was constructed using saturation mutagenesis as previously described (Qi *et al.* 2014). In brief, the entire region was divided into five sub-libraries each containing 17-18 amino acids (~500 mutants in each sub-library). NNK (N: A/T/C/G, K: T/G) was used to replace each amino acid. The oligos, each of which contains one random codon, were synthesized by IDT. The mutated region was ligated to the flanking constant regions, subcloned into the pFNX-HCV plasmid and then transformed into bacteria. The pFNX-HCV plasmid carrying the viral genome was synthesized in Dr. Ren Sun’s lab based on the chimeric sequence of genotype 2a HCV strains J6/JFH1.

### Cell culture

The human hepatoma cell line (Huh-7.5.1) was provided by Dr. Francis Chisari from the Scripps Research Institute, La Jolla. The cells were cultured in T-75 tissue culture flasks (Genesee Scientific) at 37 ^o^C with 5% CO**2**. The complete growth medium contained Dulbecco’s Modified Eagle’s Medium (Corning Cellgro), 10% heat-inactivated Fetal Bovine Serum (Omega Scientific), 10 mM HEPES (Life Technologies), 1x MEM Non-Essential Amino Acids Solution (Life Technologies) and 1x Penicillin-Streptomycin-Glutamine (Life Technologies).

### Selection of mutant viruses

Plasmid mutant library was transcribed *in vitro* using T7 RiboMAX Express Large Scale RNA Production System (Promega) and purified by PureLink RNA Mini Kit (Life Technologies). 10 μg of *in vitro* transcribed RNA was used to transfect 4 million Huh-7.5.1 cells via electroporation by Bio-Rad Gene Pulser (246 V, 950 μF). The supernatant was collected 6 days post transfection and virus titer was determined by immunofluorescence assay. The viruses collected after transfection were used to infect ~2 million Huh-7.5.1 cells with an MOI at around 0.1-0.2. The five sub-libraries were passaged for selection separately. For the three different levels of selection pressure, the growth media was supplemented with 10 pM, 40 pM and 100 pM HCV NS5A inhibitor Daclatasvir (BMS-790052), respectively. The supernatant was collected at 6 days post infection.

### Preparation of Illumina sequencing samples

For each sample, viral RNA was extracted from 700 μl supernatant collected after transfection and after selection using QIAamp Viral RNA Mini Kit (Qiagen). Extracted RNA was reverse transcribed into cDNA by SuperScript III Reverse Transcriptase Kit (Life Technologies). The targeted region in NS5A (51-54 nt) was PCR amplified using KOD Hot Start DNA polymerase (Novagen). The Eppendorf thermocycler was set as following: 2 min at 95 °C; 25 to 35 three-step cycles of 20 s at 95 °C,15 s at 52-56 °C (sub-library #1, 52 °C; #2, 52 °C; #3, 52 °C; #4, 56 °C; #5, 54 °C) and 25s at 68 °C; 1 min at 68 °C. The number of PCR cycles are chosen based on the copy number of cDNA templates as determined by qPCR (Bio-Rad). The PCR primers are listed in Supplementary Table 6. The PCR products were purified using PureLink PCR Purification Kit (Life Technologies) and prepared for Illumina HiSeq 2000 sequencing (paired-end 100 bp) following 5’-phosphorylation using T4 Polynucleotide Kinase (New England BioLabs), 3’ dA-tailing using dA-tailing module (New England BioLabs), and TA ligation of the adapter using T4 DNA ligase (Life Technologies). Each sample was tagged with a unique 3-bp customized barcodes, which were part of the adapter sequence and were sequenced as the first three nucleotides in both the forward and reverse reads (Wu *et al.* 2015) (Supplementary Table 7).

### Analysis of Illumina sequencing data

The sequencing data were parsed by SeqIO function of BioPython. The reads from different samples were de-multiplexed by the barcodes and mapped to the entire mutated region in NS5A by allowing at maximum 5 mismatches with the reference genome (Supplementary Data 3) (Qi *et al.* 2014). Since both forward and reverse reads cover the whole amplicon, we used paired reads to correct for sequencing errors. A mutation was called only if it was observed in both reads and the quality score at the corresponding position was at least 30. Sequencing reads containing mutations not supposed to appear in our single-codon mutant library were excluded from downstream analysis. The sequencing depth for each sub-library is at least ~10^5^ and two orders of magnitude higher than the library complexity.

### Calculation of relative fitness

For each condition of selection experiments (i.e. different concentration of Daclatasvir [DCV]), the relative fitness (RF) of a mutant virus to the wild-type virus is calculated by the relative changes in frequency after selection,

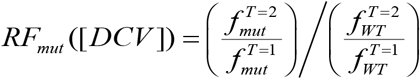

where 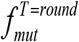 and 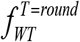 is the frequency of the mutant virus and the wild-type virus at round 1 (after transfection) or round 2 (after infection). The fitness of wild-type virus is normalized to 1. The fitness values estimated from one round (round 1 to round 2) have been shown to be highly consistent to estimated based round 0 to round 1 (Supplementary Figure 2), and estimates from multiple rounds of selection (Qi *et al.* 2014). A mutant was labeled as “missing” if the mutant’s frequency in the plasmid library was less than 0.0005 (RF=NaN, see Supplementary Data 1 and 2). A mutant was labeled as “lethal” if the mutant’s frequency after transfection was less than 0.0005, or its frequency after infection was 0 (RF=0) (Qi *et al.* 2014).

The selection coefficient is defined in the context of discrete generations (Chevin 2010)

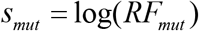

The threshold for beneficial mutations is chosen as 2*σ_silent_*, where *σ_silent_* is the standard deviation of the selection coefficients of synonymous mutations (Figure 1). The fitness effects of non-synonymous mutations leading to the same amino acid substitution were averaged to estimate the fitness effect of the given single amino acid substitution.

### Fitting the distribution of fitness effects of beneficial mutations

The distribution of selection coefficients of beneficial mutations were fitted to a Generalized Pareto Distribution following a maximum likelihood approach (Beisel *et al.* 2007),

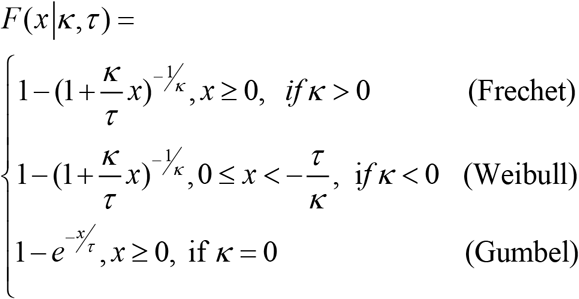

Only mutations with selection coefficients higher than the beneficial threshold 2*σ_silent_* were included in the distribution of beneficial mutations. The selection coefficients were normalized to the beneficial threshold. The shape parameter κ determines the tail behavior of the distribution, which can be divided into three domains of attraction: Gumbel domain (exponential tail, κ = 0), Weibull domain (truncated tail, κ < 0) and Fréchet domain (heavy tail, κ > 0). For each selection condition, a likelihood ratio test is performed to evaluate whether the null hypothesis κ = 0 (exponential distribution) can be rejected.

### Fitting the distribution of fitness effects to Fisher’s Geometrical model

Fisher’s Geometrical Model predicts that the distribution of fitness effects of mutations is distributed according to a negative displaced gamma distribution (Martin and Lenormand 2006a, Bank et al. 2014). This distribution has a shape parameter (α), a scale parameter (β), and a displacement parameter (so). We assume that selection coefficients are measured with a normally distributed measurement error with standard deviation *σ_s_nent·* Thus, the observed distribution of selection coefficients is modeled as the sum of a gamma and normally distributed random variable. We use the NormalGamma package in R to numerically compute the normal-gamma density function (Plancade *et al.* 2012). Maximum likelihood estimates of the parameters of the negative displaced gamma distribution are obtained with L-BFGS-B optimization implemented in the R function optim.

### Inferring drug resistance from fitness data

We can quantify the drug resistance of each mutant in the library by computing its fold change in relative fitness,

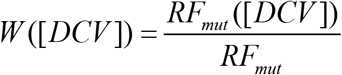

Here *RF_mut_* is the relative fitness of a mutant under the natural condition (i.e. no drug). W is the fold change in relative fitness and represents the level of drug resistance relative to the wild type. W > 1 indicates drug resistance, and W < 1 indicates drug sensitivity.

This empirical measure of drug resistance can be directly linked to a simple pharmacodynamics model (Rosenbloom *et al.* 2012), where the viral replicative fitness is modeled as a function of drug dose,

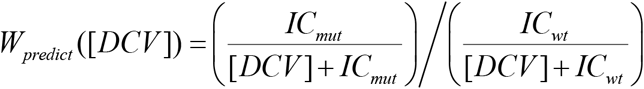

Here IC denotes the half-inhibitory concentration. The Hill coefficient describing the sigmoidal shape of the dose response curve is fixed to 1, as used in fitting the dose response curves of wild-type virus and validated mutant viruses (Supplementary Figure 5). The drug resistance score *W* inferred from fitness data is consistent with the drug resistance score *W_predict_* predicted from dose response curves of validated mutants (Supplementary Figure 6).

### Calculation of relative solvent accessibility

DSSP (http://www.cmbi.ru.nl/dssp.html) was used to compute the Solvent Accessible Surface Area (SASA) (Kabsch and Sander 1983) from the HCV NS5A protein structure (PDB: 3FQM) (Love *et al.* 2009). SASA was then normalized to Relative Solvent Accessibility (RSA) using the empirical scale reported in (Tien *et al.* 2013).

### Predictions of protein stability

ΔΔG (in Rosetta Energy Unit) of HCV NS5A mutants was predicted by PyRosetta (version: “monolith.ubuntu.release-104”) as the difference in scores between the monomer structure of mutants (single amino acid mutations from site 32 to 103) and the reference (PDB: 3FQM). The score is designed to capture the change in thermodynamic stability caused by the mutation (ΔΔG) (Das and Baker 2008). The reference sequence of NS5A in the PDB file (PDB: 3FQM) is different from the WT sequence in our experiment by 20 amino acid substitutions. Thus instead of directly comparing ΔΔG to fitness effects of individual mutations, we used the median ΔΔG caused by amino acid substitutions at each site.

The PDB file of NS5A dimer was cleaned and trimmed to a monomer (chain A). Next, all side chains were repacked (sampling from the 2010 Dunbrack rotamer library (Shapovalov and Dunbrack 2011)) and minimized for the reference structure using the talaris2014 scoring function. After an amino acid mutation was introduced, the mutated residue was repacked, followed by quasi-Newton minimization of the backbone and all side chains (algorithm: “lbfgs_armijo_nonmonotone”). This procedure was performed 50 times, and the predicted ΔG of a mutant structure is the average of the three lowest scoring structures.

We note that predictions based on NS5A monomer structure were only meant to provide a crude profile of how mutations at each site may impact protein stability. Potential structural constraints at the dimer interface have been ignored, which is further complicated by the observations of two different NS5A dimer structures (Tellinghuisen *et al.* 2005; Love *et al.* 2009).

### Diversity of HCV sequences identified in patients

Aligned nucleotide sequences of HCV NS5A protein were downloaded from Los Alamos National Lab database (Kuiken *et al.* 2005) (all HCV genotypes, ~2600 sequences total) and clipped to the region of interest (amino acid 18-103 of NS5A). Sequences that caused gaps in the alignment of H77 reference genome were manually removed. After translation to amino acid sequences, sequences with ambiguous amino acids were removed (~2300 amino acid sequences after filtering). The sequence diversity at each amino acid site was quantified by Shannon entropy.

### Data and reagent availability

All research materials are available upon request. Raw sequencing data have been submitted to the NIH Short Read Archive (SRA) under accession number: BioProject PRJNA395730. All scripts have been deposited to https://github.com/leidai-evolution/DFE-HCV.

### Ethics Statement

The use of human cell lines and infectious agents in this paper is approved by Institutional Biosafety Committee at University of California, Los Angeles (IBC #40.10.2-f).

## Acknowledgements

We thank Daniel Weinreich and two anonymous reviewers for constructive comments on the manuscript. L.D. was supported by HHMI Postdoctoral Fellowship from Jane Coffin Childs Memorial Fund for Medical Research. N.C.W. was supported by Croucher Foundation Fellowship. R.S. was supported by NSFC 81172314, NIH DE023591 and NIH CA177322.

## Author contributions

L.D., Y.D., H.Q. and R.S. designed the experiments. L.D., H.Q. and Y.D. performed the experiments. L.D. and Y.D. analyzed the experimental data. L.D., E.W. and Y.D. performed the bioinformatics analyses. C.D.H. and L.D. performed the analysis on FGM. L.D. wrote the first draft of the manuscript, with revisions from Y.D., H.Q., N.C.W., J.O.L-S., and R.S. All authors discussed the results and commented on the manuscript.

